# DNA, Morphology, and Ecology Resurrect Previously Synonymized Species of North American *Stereum* and Suggest Extensive Undescribed Global Diversity

**DOI:** 10.1101/2020.10.16.342840

**Authors:** Sarah DeLong-Duhon, Robin K. Bagley, Andrew A. Forbes

## Abstract

*Stereum* is an exceedingly common but taxonomically confounding genus of basidiomycete fungus with a cosmopolitan distribution. Lack of consensus about morphological and geographic boundaries of many *Stereum* species has resulted in a lack of consistency in identification of physical specimens, a problem that cascades to their associated published DNA sequences. A critical initial step towards addressing these issues is determining the scope of the problem. Here, we first use integrative taxonomy to delimit species in the North American *Stereum ostrea* complex. We use morphological and ecological characters, alongside ITS rDNA sequences of specimens from midwestern and eastern North America to show that “*Stereum ostrea”* in this region is a complex of at least three reproductively isolated sister species: *S. lobatum*, *S. fasciatum*, and *S. subtomentosum*. We then extend lessons from this case study to a set of publicly available *Stereum* ITS sequences to assess the accuracy of species names represented by existing sequence data. ASAP species delimitation successfully discriminates among the three newly revealed species in the *S. ostrea* species complex, but also reveals considerable cryptic diversity across global *Stereum* and widespread inconsistency in application of species names. Though ITS alone should not be used to delimit species or describe evolutionary relationships, its application here helps direct new hypotheses and suggests several areas of *Stereum* taxonomy that require revision. The critical future work of disentangling *Stereum* taxonomy and evolution should combine a multilocus genetic approach with morphology, ecology, and a global sampling strategy.

---

The most recent estimate of global fungal diversity predicts that there are 2.2 to 3.8 million species of fungi, but only ~120,000 have been formally described, and fewer than 23,000 have ITS sequences available on NCBI GenBank (Hawksworth and Lücking 2017; Vu et al. 2014). New species of fungi are often discovered as cryptic lineages previously lumped under single, well-established names (e.g., for *Amanita muscaria* and *Cantharellus cibarius* species complexes; Geml et al. 2006; Buyck and Hofstetter 2011). While the members of some species complexes may initially be difficult to separate morphologically, features useful in discriminating among species can become clear after phylogenetic analysis and trait-mapping (e.g., Peintner et al. 2019). Critically, the differences among cryptic species can be economically relevant, such as with the fungal plant pathogen *Magnaporthe grisea* and the morphologically indistinguishable *M. oryzae*, which infect crabgrass – a common invader of residential lawns – and rice, respectively (Couch and Kohn 2002).

A second complication presented by incomplete understanding of species boundaries among fungi is the lost opportunity to study the circumstances underlying their diversity. Fungi are not only incredibly speciose, but also morphologically and ecologically diverse, with similar species often occupying different habitats/substrates (Seitzman et al. 2011, Wibberg et al. 2021). In this way, they present fantastic systems to explore how divergent selection along a variety of ecological axes might influence their biodiversity. However, given the absence of robust taxonomic resources and a subsequent lack of understanding of where one species begins and another ends, questions about fungal evolution are all but impenetrable.

One understudied fungal group that may harbor many undiscovered species and that may be a good candidate for future ecological speciation research is *Stereum*, a diverse genus of shelf-like wood-decay fungi in the Russulales order that is common in wooded biomes throughout the world. Though *Stereum* has been the focus of extensive bioprospecting (Doljak et al. 2006; Hybelhauerová et al. 2008; Tian et al. 2020), and some economically important species are relatively well-researched (Stenlid and Vasiliauskas 1998; Čermák et al. 2004), the below-genus level taxonomy of *Stereum* has not been subjected to rigorous phylogenetic analysis. This disconnect between the slow progress toward molecular taxonomy in *Stereum* and interest in the potentially useful properties of specific *Stereum* species can be problematic, especially where species descriptions include uninformative or deceptive morphological characters that may lead to misidentification of study specimens.

*Stereum ostrea* is an exemplar of a species with a fraught taxonomic history (Lloyd 1913). While the name *S. ostrea* has been applied to collections around the world, it is unlikely that these varied collections belong to a single phylogenetic species given pre-Anthropocene obstacles to dispersal. First used to describe a collection from the island of Java in Indonesia, the name *S. ostrea* has been consistently applied to specimens from North America since a publication by Lentz (1955) placed it in synonymy with two other putative species, *S. fasciatum* and *S. lobatum*, with which Welden (1971) and Chamuris (1985, 1988) concurred. Even before this, *S. ostrea* was considered either a synonym of *S. fasciatum* (Burt 1920; Pilát 1930; Banerjee 1935; Hendrickx 1948) or of *S. lobatum* (Massee 1890; Cooke 1892; Höhnel and Litschauer 1907; Reinking 1920; Boedijn 1940). Demoulin (1985) argued in favor of *S. fasciatum* and *S. lobatum* being two species, distinct from one another and from *S. ostrea*, but the literature remained divided (Eicker and Louw 1998). Currently, *S. fasciatum* is considered a synonym of *S. ostrea* (Species Fungorum, February 2022, http://www.speciesfungorum.org/), and despite being an accepted name, *S. lobatum* is rarely used. Another similar species, *S. subtomentosum,* has both been confused with *S. fasciatum* and proposed as belonging to a species complex with *S. hirsutum* (Pouzar 1964; Welden 1971; Chamuris 1988; Ginns and Lefebvre 1993).

We sought to 1) test the hypothesis that “*S. ostrea*” in North America is not a single species but a complex of several species and 2) to use thresholds of sequence divergence shown to separate putative species in this case study of *S. ostrea*, with the overall goal of interrogating the potential scale of unrealized global *Stereum* diversity. We collected >50 *Stereum* specimens from eastern North America, documented their morphology and ecology, and collected nuclear rDNA ITS1-5.8S-ITS2 (internal transcribed spacer barcode) sequence data. We also used our sequence data, 13 sequences from the New Zealand Fungarium (PDD), and 415 ITS sequences tagged as *Stereum* that we harvested from online sequence repositories, for a preliminary analysis of global cryptic species diversity in *Stereum*. Overall, we find support for multiple species within *S. ostrea* as well as evidence of rampant misidentification of species within global datasets. These findings are a first step toward creating a robust global phylogeny for the genus *Stereum*.

## Materials and Methods

### Collection and identification

We collected 36 *Stereum* basidiocarps from sites in midwestern and eastern North America, with the most intensive sampling in Iowa and some supplemental collections made in Alabama and Florida (Supplementary Table 1). For identification we used the dichotomous key and morphological descriptions from Chamuris (1988), the most comprehensive recent publication on *Stereum*. Based on morphology, we initially identified our collections as *S. ostrea*, *S. hirsutum*, *S. complicatum*, *S. gausapatum*, *S. sanguinolentum*, and *S. striatum*. We used iNaturalist to record photographs, dates of collection, and approximate GPS coordinates of collection location (Supplementary Table 1). We also recorded the substrate from which samples were collected, whether substrate was hardwood or conifer, and if the hymenium of the mushrooms changed color when bruised. We air dried all collections and preserved samples in polyethylene plastic bags.

### High Resolution Imaging

We photographed exemplar *Stereum* with a Canon EOS 60D camera and a Canon MP-E 65mm macro lens (Canon USA, Melville, NY) mounted on a StackShot automated macro rail (Cognisys Inc., Traverse City, MI). Images were processed in Zerene Stacker software using PMax method (Zerene Systems LLC, Richland, WA).

### DNA extraction, PCR and sequencing

We used a CTAB and liquid nitrogen method adapted from Chen et al. (2010) to extract DNA from 3 x 3 mm pieces of basidiocarp from each collection, taking care to exclude as much dirt and debris as possible. We used DNA diluted 1:20 with molecular grade water for PCR amplification using ITS1‐F (CTTGGTCATTTAGAGGAAGTAA) and ITS4 (TCCTCCGCTTATTGATATGC) primers with the following thermocycler program: 3 min @ 94 C, 25 cycles (30 s @ 94 C, 30 s @ 55 C, 30 s @ 72 C), 2 min @ 72 C (White et al. 1990; Gardes and Bruns 1993). We cleaned PCR products with Exo-SAP following manufacturer protocols, then sequenced PCR products for both forward and reverse directions on an Applied Biosystems ABI 3730 DNA Analyzer (Thermo Fisher Scientific, Massachusetts) housed in the Roy J. Carver Center for Genomics in the University of Iowa Biology Department.

### Phylogenetic analysis

We used Geneious 8.1.7 (http://www.geneious.com/) for alignment of forward and reverse sequences for each collection, which we then manually checked and trimmed. We chose *Xylobolus subpileatus*, another member of the Stereaceae, as an outgroup. This specimen was collected in Florida by S.G.D. and sequenced by the Smith Lab at the University of Florida. We also obtained North American and Eurasian *S. hirsutum* sequences from Barcode of Life Database (BOLD) and GenBank. We used MAFFT (Katoh et al. 2002) via the CIPRES server (Miller et al. 2010) to align chosen sequences, then generated a maximum likelihood tree using RAxML (Stamatakis 2014) via CIPRES. For computation of the Bayesian tree in Supplementary Figure 1, we used MrBayes 3.2.7 (Ronquist et al. 2012) with a GTR+I+Γ substitution model for 200,000 generations, with sampling every 100 generations. Sequences generated during this study were deposited in the GenBank database (Supplementary Table 1). Alignment and tree files were deposited to Treebase (Submission ID: 27646).

### ASAP Species Delimitation

We downloaded all available sequences from GenBank on October 18^th^, 2021, using the search parameters “Stereum“[Organism] AND “internal transcribed spacer“[All Fields], for a total of 547 sequences. We also retrieved 13 *Stereum* sequences from BOLD, used 13 sequences from the New Zealand Fungarium (PDD) courtesy of Dr. Jerry Cooper, and included 32 of the 36 *Stereum* sequences we generated for this study (Supplementary Table 2). Four of our sequences were not used due to inadequate sequence length.

After an initial alignment using MAFFT we discarded sequences that were a) not *Stereum*, b) of low-quality, and c) were missing more than ~30bp of the final trimmed sequence length of the alignment. Examples of sequences found to not to be *Stereum* (16 total) were *Trametes hirsuta*, *Penicillium*, *Scytinostroma*, and *Khuskia*, according to the closest GenBank BLAST matches. Sequences were presumed to be of low quality where there were significant random-seeming changes in the highly conserved 5.8S region or other smaller regions that are conserved among all other *Stereum* ITS sequences. Sequences were pared down to ensure no large gaps on either end of the alignment to increase accuracy of Assemble Species by Automatic Partitioning (ASAP), and sequences missing more than 30bp on either end of the final alignment were discarded (67 total). In total, these filters removed ~100 specimens. We realigned the remaining 460 sequences with MAFFT and ran an ASAP analysis via the web portal (https://bioinfo.mnhn.fr/abi/public/asap/) with default parameters: substitution model Jukes-Cantor (JC69) (Puillandre et al. 2021). Output files were downloaded and used to create Figure 4. Metadata for sequences in the final alignment can be found in Supplementary Table 2. To create a preliminary sense of how putative *Stereum* species might be related, we inferred a neighbor-joining phylogeny using a Jukes-Cantor distance model and aligned ASAP species groupings with its tips.

## Results

For our new North American *Stereum* collections, maximum likelihood (Fig. 1) and Bayesian trees (Supplementary Fig. 1) were largely in agreement, with the sole exception being the placement of the single *Stereum striatum* sequence. Both trees show *Stereum* sorting out into several clades. Specimens initially identified as *S. ostrea* formed one large monophyletic group with three individuals initially identified as *S. hirsutum*, but this was split further into three smaller, well-differentiated clades (Fig. 1). Sequences within each clade differed from one another by 1–3%, while sequences among the three clades differed from one another by 7–14%.

**Figure 1.**
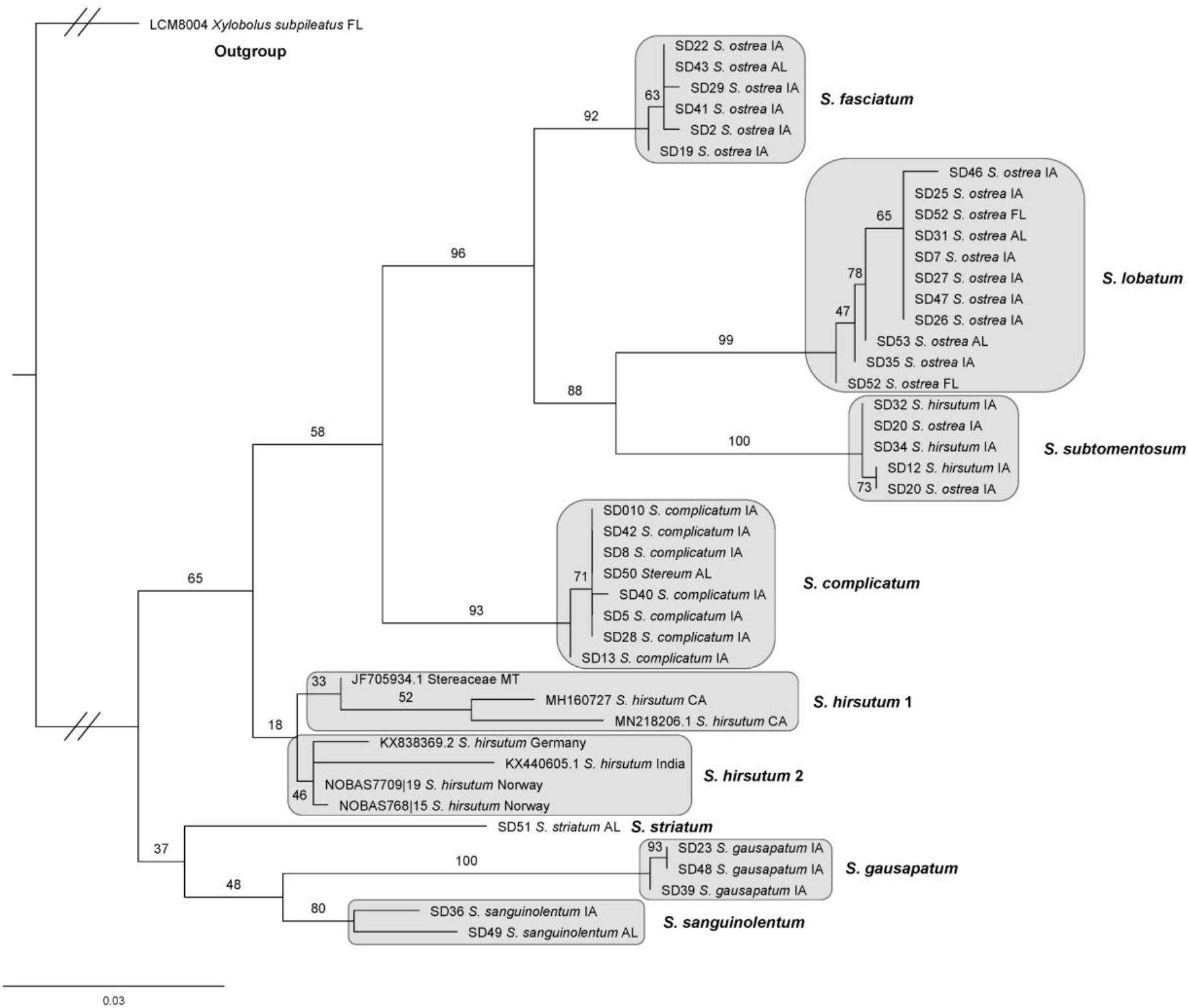
Maximum likelihood phylogeny of *Stereum* collections generated from ITS sequence data. Bootstrap values are above branches. Scale bar represents the number of nucleotide changes per site. The topmost three clades show strong support, suggesting three distinct members of the *Stereum ostrea* species complex in midwestern and eastern North America, which we identify here as *S. fasciatum*, *S. lobatum*, and *S. subtomentosum*, based on morphological differences (Fig. 2 & 3).

**Figure 2.**
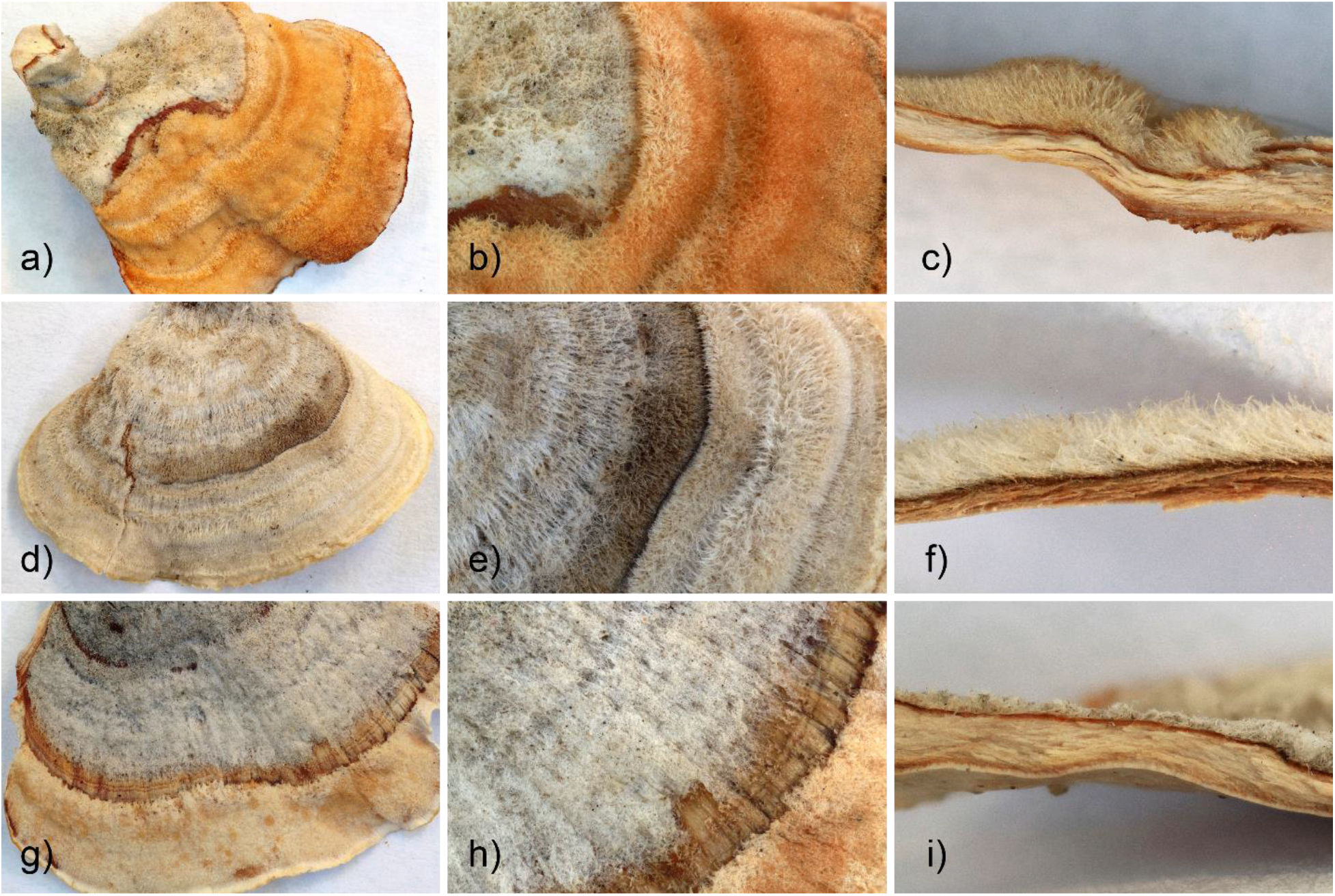
Morphological comparison of a-c) *Stereum subtomentosum*, d-f) *S. fasciatum*, g-i) *S. lobatum*. Respectively, the columns show wide views of the basidiocarps, magnified views of the tomentum texture, and cross sections. *S. subtomentosum* tomentum is long, wooly and tufted, *S. fasciatum* tomentum is coarse and clumped in tufts, and *S. lobatum* tomentum is short, matted, and felted. Note that cap hair is best viewed under magnification.

**Figure 3.**
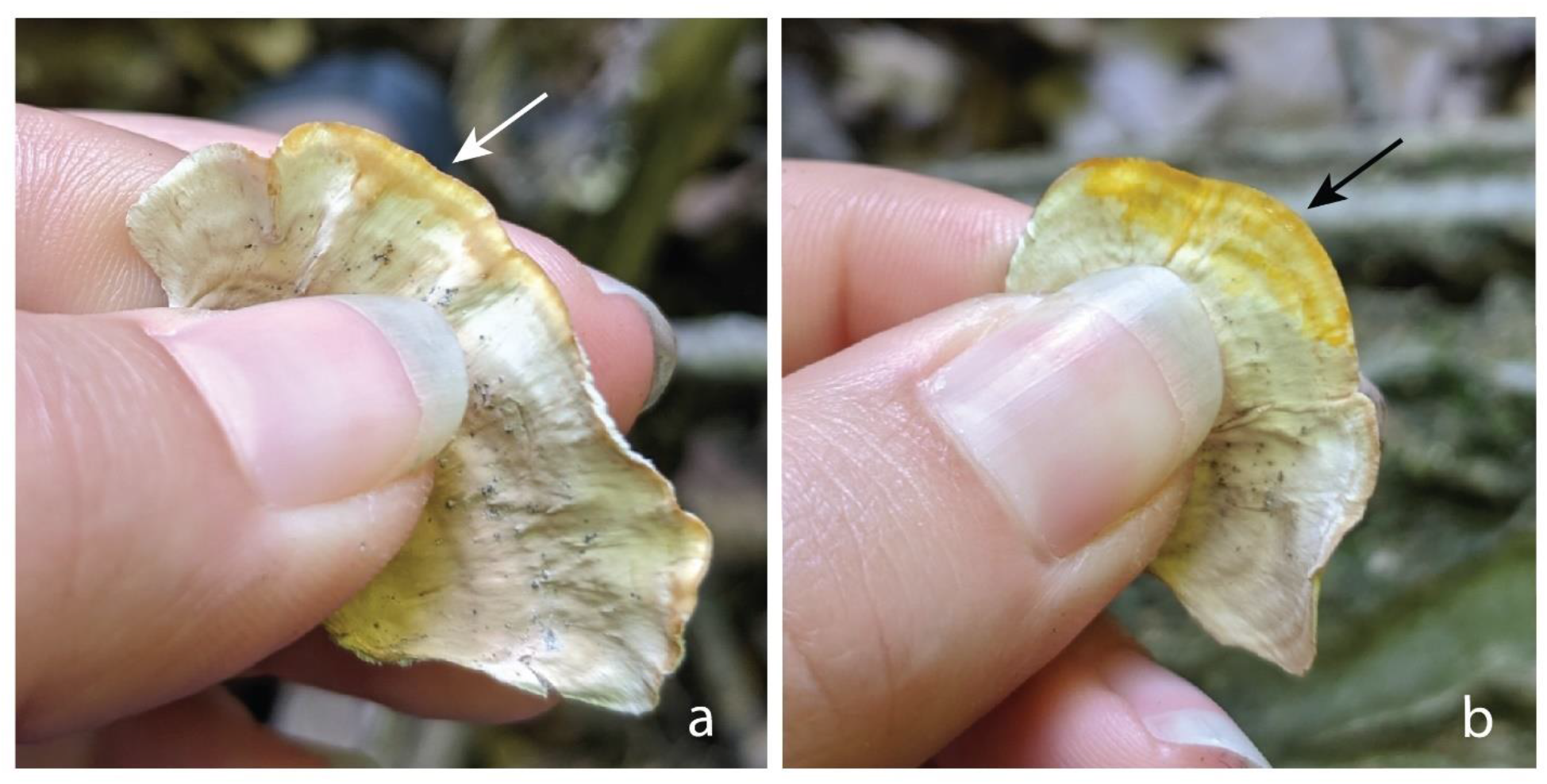
Comparison of color staining between a) *S. fasciatum* and b) *S. lobatum*, which show slight darkening, and bright yellow staining, respectively. *S. subtomentosum* (not shown) exhibits the same bright yellow staining as *S. lobatum*. Note that both specimens were dry, and re-wetted to investigate color staining.

The specimens in the first of the three subclades (Fig. 1) all share features of the previously synonymized species *S. fasciatum*, as defined by many authors including Burt (1920) and Demoulin (1985). These specimens feature a cap clothed in coarse hair that is resistant to wearing off in bands, and with these hairs gathered in individual clumps that are best observed with a hand lens or dissecting microscope (Fig. 2a–c). If bruised (when basidiocarps are fresh) or wetted (when basidiocarps are dry) the smooth hymenium (undersurface) appears wet but does not stain color (Fig. 3a).

Similarly, the specimens in the second clade (Fig. 1) showed features consistent with the previously synonymized *S. lobatum* (Burt 1920; Demoulin 1985). The cap is clothed in matted, felted hairs that quickly begin wearing off, from the edge of the cap inwards, in concentric bands exposing the shiny chestnut-brown context beneath (Fig. 2d–f). Importantly, the short cap hairs are matted and tangled together in such a way that individual hairs are very difficult to observe. When bruised or wetted the hymenium stains a bright yellow color (Fig. 3b).

Specimens in the third clade shared some features with both *S. fasciatum* and *S. lobatum*, however the basidiocarps were usually thicker, more irregular (i.e., growth zones not evenly concentric, caps radially folded and overlapping), and both cap and hymenium often a richer coffee-brown color. The cap is covered in clumping hairs that are typically longer and woolier than *S. fasciatum* and wear off in bands more readily but not to the extent of *S. lobatum* (Fig. 2g–i). When bruised or wetted the hymenium stains a bright yellow color like *S. lobatum* (Fig. 2b). Overall, the morphology agrees with descriptions of *S. subtomentosum* (Pouzar 1964; Chamuris 1988).

Specimens identified before sequencing as belonging to other species of *Stereum*, including *S. complicatum*, *S. gausapatum*, and *S. sanguinolentum*, formed reciprocally monophyletic clades with ITS sequences differing within clades by 1–5% (Fig. 1). The single sequence of *S. striatum* was placed differently in the two trees but differed from all other sequences by more than 7%. *S. hirsutum* sequences acquired from GenBank and BOLD databases differed from one another by 1–6%, but only as much as 4% within the two geographically isolated (North America vs. Eurasia) clades.

ASAP species delimitation results for the 415 sequences downloaded from databases, plus our own samples and those from PDD, suggest that the full dataset contains between 19 to 74 species, with the best scoring estimates being 19, 20, and 30 (ASAP scores 3.5, 3.5, and 4.0, respectively). Figure 4 includes two of the three estimates with the lowest (best) ASAP scores. The estimate of 19 was omitted in the final figure due to its similarity to the 20 species estimate. In most cases, sequences that ASAP assigned to the same putative species were associated with the names of >1 *Stereum* species. Additionally, one *Stereum* species name was often assigned to two or more putative species. For example, at an ASAP score of 4, the name *S. sanguinolentum* is found in 13 different assignations, differing in percentage sequence similarity by as much as 13%. Additionally, *S. hirsutum* was found in 9 different assignations, and the name *S. ostrea* (including *S. cf. ostrea*) was found in 6 different assignations.

**Figure 4.**
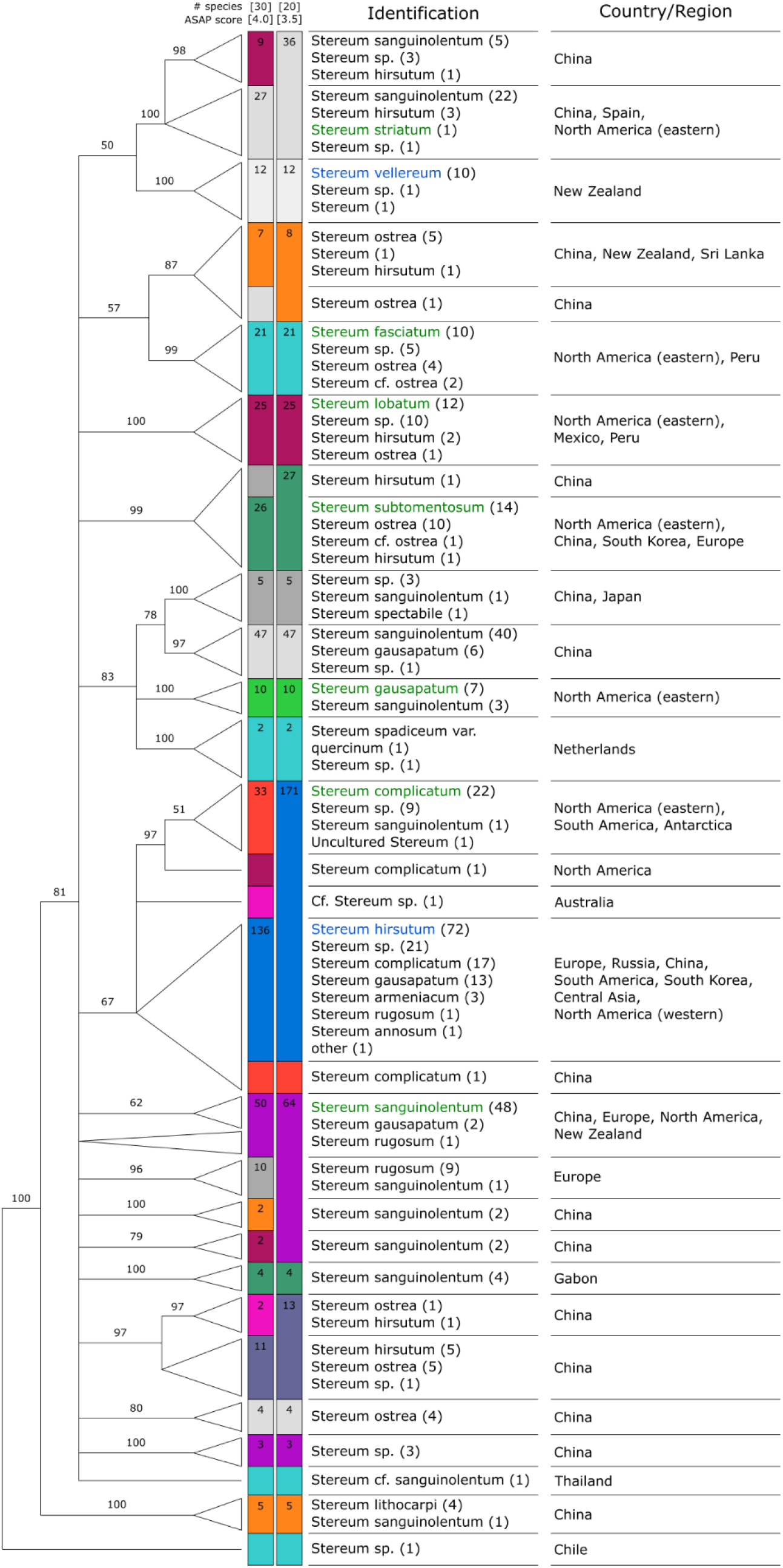
ASAP species delimitation results for 460 *Stereum* ITS sequences (see Supplementary Figure 2 for full results). Best scoring estimates – which also sorted our own collections from Figure 1 correctly by species – resulted in estimates of 20 (ASAP score = 3.5) or 30 (ASAP score = 4.0) species, depicted by the colored boxes in either column. Numbers in each box represent the number of sequences for each named species (one if blank). Box size does not accord with sequence number. Identification and Country/Region data summarizes metadata for each species. Names highlighted in green represent our own collections reported in Figure 1, and names highlighted in blue represent names confirmed with photographic evidence. The dendrogram to the left is provided only as a preliminary estimate of relationships among ITS sequences.

## Discussion

Our results support the hypothesis that the North American fungi that have been colloquially lumped under the name “*Stereum ostrea*” are actually at least three reproductively isolated and morphologically distinguishable species: *S. fasciatum*, *S. lobatum*, and *S. subtomentosum*. The inference of reproductive isolation is based on the observation that while all three species are often found growing at the same site on the same substrate, there is strong concordance between ITS sequence identity and morphology, such that they appear not to be hybridizing despite often living in sympatry. Demoulin (1985) argued that *S. fasciatum* and *S. lobatum* were distinct species based on differing morphology and lack of fusion while growing on the same substrate, and our results support this claim with new genetic evidence.

As a consequence of these new data, these three *Stereum ostrea*-like species should be again recognized as legitimate and distinct species, and we recommend that the name *S. ostrea* no longer be used for North American collections. These three species in the midwestern and eastern North America can be reliably differentiated by a combination of three features, 1) texture of cap hairs, 2) presence and extent of concentric banding, and 3) color staining of the hymenium when bruised or wetted (Figs 2–3). Other traits such as shape and color are often too variable to be used as distinguishing characteristics on their own and should only be used to inform an ID in conjunction with the aforementioned features. Our Figures 2 and 3 along with images for each specimen on iNaturalist (Supplementary Table 1) are provided to assist future naturalists and researchers in distinguishing among these species.

Support for *S. ostrea* being a complex of species different in both morphology and ITS sequence endorses them as a within-genus standard for using ITS sequence divergence to develop preliminary species hypotheses for *Stereum*. ASAP clearly delineates *S. fasciatum*, *S. lobatum*, and *S. subtomentosum* from one another at both score levels in Figure 4, and so we suggest – at least as an initial hypothesis – that many other divisions at these same levels may also represent reproductively isolated species. If this is true even in part, the sharing of names across many putative species designations and the clustering of different names into the same designation must represent misidentification, inadequacies in the taxonomy, or more likely, both.

Both levels of ASAP scores maintained separation among the species treated in our North American collections, and added several sequences to each clade, many of them with different assigned names (Fig. 4). All putative *S. fasciatum* and *S. lobatum* sequences were also from collections made in the Americas, while sequences aligning with our *S. subtomentosum* clade were from collections in North America, Europe, and East Asia. *S. complicatum* sequences were from both South and North America with the exception of one enigmatic sequence from a cave in Antarctica (KC785597).

Sequencing all *Stereum* more thoroughly on a global scale will be necessary to elucidate species’ geographic distributions. The current spotty records likely underlie, for instance, a lack of sequence representatives for some species from their country of typification - such as *S. gausapatum* which is described from France but is only represented in our analysis by sequences from eastern North America. Species names may also be historically applied to the wrong taxon, so if for example we later find that *S. gausapatum* in France is genetically distinct from North American collections of the same name, an alternate name would need to be found. Additionally, there is high ITS sequence diversity – as much as 8.2% difference within-species – among putative *S. hirsutum* based on the 30 species ASAP estimate. In the two highest species estimates, *S. hirsutum* is divided into smaller groups that correlate with geographic origin, with all sequences from China and North America in their own group. This suggests the presence of cryptic species. *Stereum hirsutum* is known for its morphological variability, which has led to the typification of multiple varieties and forms (Turam et al. 2008). Further study of these *S. hirsutum* groups may help determine whether *S. hirsutum* is one globally widespread species, or instead a complex of morphologically similar species.

Importantly, ASAP and similar methods should not be used alone to make definitive conclusions about species limits, but rather as a first step informing future taxonomic work. Though we argue – based on having ground-truthed ASAP scores with our own data – that many of these putative species may be isolated from one another, we also see evidence that ASAP can be prone to error. Only one of the three highest scoring partitions recognized *S. complicatum* as a species separate from *S. hirsutum*, despite the large and consistent sequence difference (4.5– 10%) and high support values for *S. complicatum* in Figure 1, and differences in geographic distribution and morphology. Though it may have problems, species delimitation using ASAP and similar approaches like ABGD and bPTP has proved useful for making species number estimates in other taxonomically tricky taxa such as Chaitophorinae aphids, physinine snails, and inquiline wasps (Zhu et al. 2017, Ward et al. 2020, Young et al. 2021). The strength of this approach lies not it its ability to finalize species boundaries, but in its ease of use, making it easily replicable with new datasets and relatively simple to interpret results. This should make it a particularly attractive option for non-professional mycologists.

A more general conclusion is that an integrative approach to *Stereum* taxonomy that includes, but is not limited to, DNA sequence data will be helpful in resolving species boundaries. In most cases, original species descriptions are only a few sentences long or quite vague. Further, *Stereum* taxonomy has a long history of synonymization such that descriptions of *Stereum* species from different authors are often conflicting, as they generalize traits of multiple species. Historically, *Stereum* has been used as a genus name to describe many corticioid fungi, many of which have been transferred to other genera both in the Stereaceae and to other genetically distant families (Larsson and Larsson 2003). Many *Stereum* are still accepted as synonyms of better-known *Stereum* species, but if our study is of any indication, some of those names may also need to be resurrected.

Our work here demonstrates a need to explore the rest of the *Stereum* genus, in order to provide taxonomic resources for those who study other aspects of *Stereum*. Their ecological roles as parasites and wood decay fungi, as well as potential use in bioremediation and biomedicine, have made them an attractive organism for research. Molecular phylogenetic approaches will be necessary as a supplemental tool to delineate *Stereum* species, as morphology alone has proved inadequate in the past, and we predict that the key features useful in differentiating the members of the North American *S. ostrea* species complex treated here (i.e., tomentum, staining, banding) may not delineate *Stereum* species so clearly elsewhere in the world. While using a single locus was sufficient for this regional preliminary study, more loci should be included for resolving the phylogeny of *Stereum*. By using a multilocus approach and worldwide sampling strategy, we can achieve greater resolution within this genus and other members of the Stereaceae.

## Acknowledgements

We thank Dr. Rosanne Healy for providing an outgroup sequence and valuable advice, and Dean Abel for his encouragement and gift of important literature on *Stereum*.

