## Supplementary Figure 1 for "DNA, Morphology, and Ecology Resurrect Previously Synonymized Species of North American *Stereum* and Suggest Extensive Undescribed Global Diversity"

**Supplementary Figure 1.** Bayesian phylogeny generated from ITS sequence data. Bootstrap values are above or below branches. Scale bar represents the number of nucleotide changes per site. The first three clades show strong support, suggesting *S. ostrea* in midwestern and eastern North America consists of three distinct species, which we identify as *S. fasciatum*, *S. lobatum*, and *S. subtomentosum*.

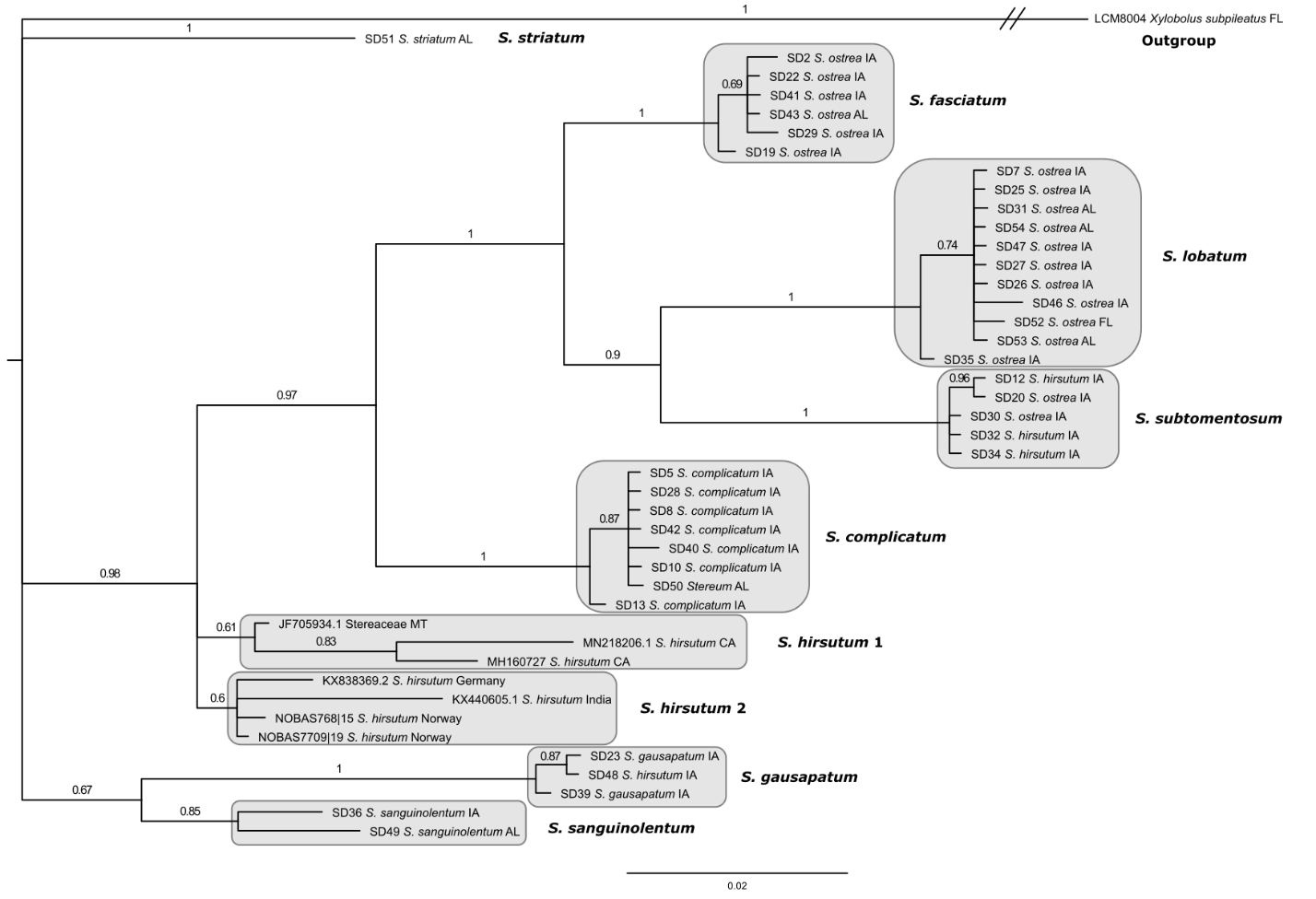
