## Supplementary figures and images for "DNA, Morphology, and Ecology Resurrect Previously Synonymized Species of North American *Stereum* and Suggest Extensive Undescribed Global Diversity"

### Supplementary Figure 2

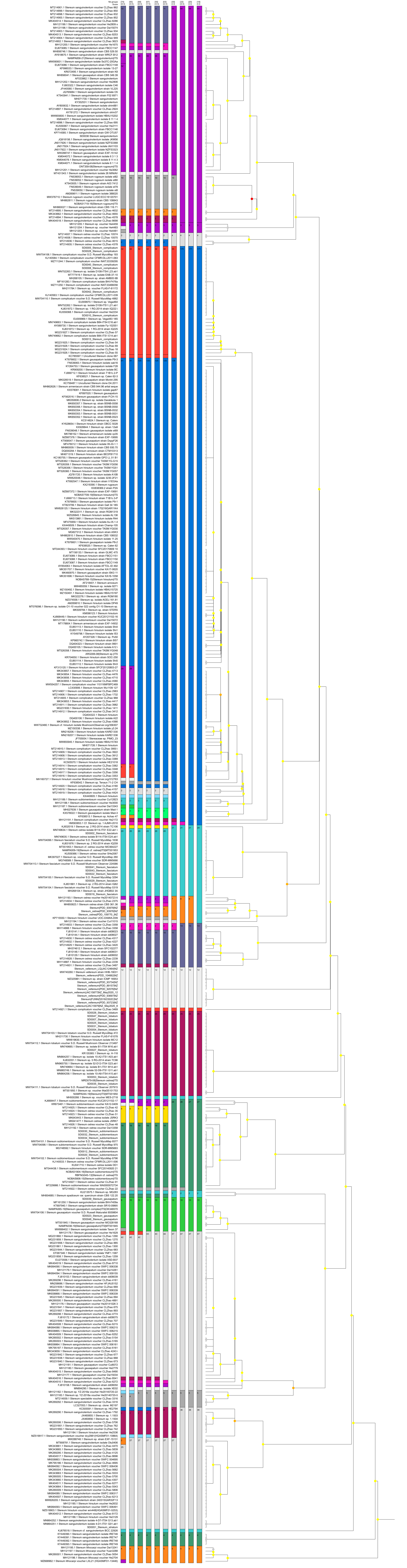
